# Systematic benchmarking of low-input whole exome sequencing workflows for longitudinal ctDNA profiling in pancreatic ductal adenocarcinoma

**DOI:** 10.64898/2026.06.29.734743

**Authors:** LGE James, GJ Thorn, C Morel, PCRFTB, HM Kocher, H Ross-Adams, C Chelala

## Abstract

Whole exome sequencing (WES) of circulating tumour DNA (ctDNA) enables longitudinal monitoring of tumour dynamics, evolution and treatment response but remains technically challenging in low-input, low-shedding settings such as pancreatic ductal adenocarcinoma (PDAC). Here, we systematically compared three commercially available low-input WES workflows incorporating Agilent (V6, V8) and Qiagen exome capture designs using ultra-low input cfDNAs extracted from multiple matched longitudinal plasma samples from PDAC patients.

Using predefined performance metrics including coverage, duplication rate and variant detection and additional metrics relevant for clinical genomic profiling in patient care, we show that all three workflows produced high-quality sequencing data, even from very low input cfDNA. Within the conditions tested here, the Agilent V8 workflow provided the most favourable balance of coverage uniformity, sequencing efficiency and hotspot coverage for low input, low tumour fraction cfDNA WES.

These findings demonstrate that workflow design, including capture footprint, substantially influences ctDNA WES performance in low-input clinical contexts. These findings are particularly relevant in early stage and/or minimal residual disease settings, where tumour fractions are low and recovery of genomic information from limited-input samples is critical.

## Background

Pancreatic ductal adenocarcinoma (PDAC) is predicted to become the second leading cause of cancer-related mortality worldwide before 2030 [1]. It has dismal 5-year survival rates of 3-12% [1, 2] largely due to late detection and few effective treatment options: ∼85% of patients are diagnosed with inoperable disease.

Despite recent efforts, scant progress has been made in using molecular biomarkers to improve patient management. Current biomarkers for assessing prognosis or monitoring treatment have limited sensitivity and/or specificity in prospective settings [3]. Blood serum carbohydrate antigen CA19-9 is the current gold-standard for PDAC, but it is seldom used alone and is uninformative in non-secretors (5-10% general population) [4]. Furthermore, PDAC displays significant inter- and intra-tumoural genetic heterogeneity together with a complex, dynamic and supportive microenvironment [5]; studies investigating how individual tumour clones evolve and respond to treatment are limited. Tissue biopsies cannot completely capture the spatial and temporal complexity of each tumour. Tracking genomic alterations would normally require multiple biopsies, which would present an additional burden on patients and is not standard care. Better tools for early disease detection, patient stratification and treatment response are thus essential to improve patient outcomes.

Liquid biopsies provide a means to identify and track a growing compendium of blood or urine-borne tumour markers like circulating tumour cells (CTCs), tumour DNA (ctDNA), miRNAs, various extracellular vesicles and tumour-educated platelets (TEPs) [6] using minimally invasive approaches. Next generation sequencing (NGS) of plasma-derived cfDNAs has identified patient-specific actionable mutations, revealed treatment response earlier than standard biomarkers or imaging, monitored clonal evolution in real time and identified the emergence of resistant clones [7–10]. However, such experiments typically rely on an abundance of plasma (≥4 ml) and input cfDNAs (≥ 10ng) to reliably detect key somatic variants and copy number alterations (CNAs) present at low variant allele frequencies (VAFs), which may not be routinely available outside a research setting [11].

cfDNA extraction and NGS capture kit technologies are rapidly improving [11], facilitating the study of tumours with lower ctDNA fractions (<1%) (e.g. PDAC, renal, bladder) [12, 13], as well as patients with early or minimal residual disease. This is particularly relevant in PDAC, where low tumour cellularity and dense stromal architecture contribute to low circulating tumour fractions, making sensitive, low-input sequencing approaches essential for clinically useful molecular profiling [6, 14].

WES of ctDNA is inherently a compromise between breadth and sensitivity. While pan-cancer gene panels offer faster, cheaper and deeper sequencing of known disease risk and/or actionable mutations, we and others have shown that WES can identify actionable patient-specific variants that would not have been detected using a gene panel capture [10, 15, 16]. WES can also capture a more comprehensive picture of tumour heterogeneity and is particularly useful for identifying complex biomarkers like tumour mutational burden (TMB), microsatellite instability (MSI) and homologous recombination deficiency (HRD) [16, 17], which have been associated with therapeutic targets in PDAC [18, 19]. However, in low-shedding tumours like PDAC, the broader scope of WES comes at the cost of reduced analytical sensitivity relative to tumour informed or highly targeted approaches.

Despite rapid advances in cfDNA sequencing technologies and exome capture designs, there has been little systematic evaluation of low-input WES workflows in longitudinal ctDNA samples from low-shedding tumours such as PDAC. This is particularly important in low-input clinical contexts, where differences in capture efficiency, molecular barcoding strategies, coverage and error profiles may have a disproportionate impact on variant detection, data quality and downstream clinical interpretation.

Here, we present a technical benchmarking study of recent low-input whole exome capture workflows (Qiagen’s QIAseq Human Exome and Agilent’s SureSelect XT Human All Exon v8) using longitudinal plasma ctDNAs samples from patients with resectable PDAC, focusing on practical performance metrics relevant to low-input clinical research settings. We further benchmark these against data generated on matched plasma samples with Agilent’s SureSelect XT2 v6.0 Human All Exon capture kits [10] as a known baseline using multiple metrics, and discuss several issues that can arise from applying such metrics to the specific case of cell-free DNA.

## Methods

We performed a systematic benchmarking of three whole-exome sequencing capture workflows using predefined performance metrics applied consistently across matched longitudinal plasma samples from PDAC patients. Longitudinal sampling enabled evaluation of workflow performance across multiple clinically distinct time points while maintaining biological context.

## Patients and samples

Whole blood, plasma and tumour biopsy tissues from patients with histologically confirmed pancreatic ductal adenocarcinoma were obtained from the Barts Pancreatic Tissue Bank (www.bartspancreastissuebank.org.uk, Research Ethics Committee reference 13/SC/0592). All patients provided written, informed consent. Two patients (P045, P095) who had undergone surgical primary tumour resection also had matched germline samples, and >3 serial plasma samples collected at clinically determined intervals following initial diagnosis or surgery. Three longitudinal plasma samples from these two patients were used for comparison: P045_A, P095_A, and P095_D, reflecting the availability of matched cfDNA material available across all capture chemistries and intended to support a controlled technical comparison of workflow behaviour rather than enabling robust statistical inference or clinical generalisation.

## Sample processing

Serial whole blood samples were collected in either RUO Cell-Free DNA Collection Tubes (Roche) or 10 ml Vacutainer K3EDTA tubes (BD Biosciences) and processed for plasma isolation within two hours of collection. Whole blood was centrifuged at 1600g for 10 minutes at room temperature using a swing-out rotor, to separate plasma from peripheral blood cells. Derived plasma and buffy coat samples were stored in 500 µl aliquots at -80 °C until extraction.

All plasma cfDNAs were extracted using QIAmp MinElute ccfDNA kit (Qiagen) immediately prior to library preparation, according to manufacturer’s instructions. For Agilent V6, cfDNA was extracted from 1.5 ml-3 ml plasma per time point. For Agilent V8 and Qiagen, cfDNA was extracted from 500 µl plasma per time point (reaction volumes adjusted for input amounts). In all cases, final extracts were eluted in 50 µl nuclease-free ultra-pure water, then re-eluted with original flow-through for a total of two elutions per extract and ∼46 µl per sample. All details per sample, per chemistry are detailed in Supplementary Table S1, including starting input amounts per reaction.

Genomic DNA was extracted using DNeasy Blood and Tissue kit (Qiagen) according to manufacturer’s instructions. Tumour tissues from P045 and P095 were macerated beforehand in pre-chilled buffer ATL with a TissueLyser (Qiagen) and 5mm stainless steel balls at 25Hz for 60s; germline DNA was extracted directly from buffy coat samples. Extracted gDNAs were sonicated to approximately 200bp (Covaris M220) and stored at -80 °C until required.

All sample quality, peak distributions and concentrations were assessed throughout using TapeStation (Agilent) and Qubit (ThermoFisher) respectively.

## Sequencing of tumour and normal DNA

Tumour and germline libraries were prepared from up to 1µg gDNA using TruSeq nano DNA kits (Illumina), according to the manufacturer’s instructions. Library preparation, whole genome sequencing and alignment was performed by Edinburgh Genomics core facility, to a mean depth of 110X (tumour) and 40X (germline) on HiSeqX (Illumina).

## Sequencing of plasma cfDNA

### Agilent V6

Plasma libraries were prepared from up to 10ng cfDNA using Rubicon ThruPLEX Plasma-Seq kits (Rubicon Genomics; Cat. R400490), as previously described (Sivapalan 2022). Exome capture was performed using SureSelect XT2 v6.0 Human All Exon (Agilent, Cat. 5190-8872) kits with i5 and i7 xGen Universal Blocking Oligos (Integrated DNA Technologies), following the manufacturer’s recommendations for compatibility with ThruPLEX libraries. Enriched libraries were quantified and equimolar amounts pooled for sequencing on HiSeq 4000 (Illumina) to 500X target depth at CRUK Cambridge Institute Genomics core facility.

### Agilent V8

Plasma cfDNA libraries were prepared using SureSelect XT HS2 DNA kits (Agilent, Cat G9985B), according to manufacturer’s instructions. Adapters including molecular barcodes which serve as unique molecular indexes (UMIs) were incorporated using 14X amplification cycles to generate dual-indexed libraries. Sample quality, peak distribution and concentration of each was determined, before normalising to the lowest concentration sample and pooling equimolar amounts of individual samples into one library. Whole exomes were captured using SureSelect XT Human All Exon v8 kits (Agilent v8, Cat. 5191-6879), with a hybridisation at 67.5°C for 21 hours, then amplified using 10X post-capture amplification cycles. Final libraries were quantified and equimolar amounts pooled for sequencing on NextSeq (Illumina) to 1290X target depth at QMUL Genome Centre.

### Qiagen Whole Exome

Plasma cfDNA sequencing libraries were prepared using QIAseq® Ultralow Input Library kit (12) (Cat. 180492), recommended for inputs 10 pg - 100 ng, with QIAseq UDI Y-adapter kit (24) (Cat. 180310) (both Qiagen) according to manufacturer’s instructions. Adapters were used diluted 1:10 in nuclease-free water, according to available sample input concentrations, with 10X amplification cycles to generate dual-indexed libraries. Sample quality, peak distribution and concentration of each was determined, before normalising to the lowest concentration sample and pooling equimolar amounts of individual samples into one library. Whole exomes were captured and amplified using QIAseq Human Exome kit (Cat. 333937) with One-4-All blocking oligos, by hybridisation at 60°C for 18 hours and 8X post-capture amplification cycles for pools <1000ng. Final libraries were quantified and equimolar amounts pooled for sequencing on NextSeq (Illumina) to ∼1128X target depth at QMUL Genome Centre.

All experimental details are summarised in Supplementary Table S2.

## Bioinformatics analysis

### Capture design and exon overlap

Manufacturer-provided BED files for the V6, V8 and Qiagen panels were analysed in Python (v3.10) using pandas [20]and PyRanges [21]. Genomic intervals were converted to PyRanges objects, and pairwise and three-way intersections were performed to identify shared regions and quantify overlapping base-pair coverage; unique regions were defined by subtracting overlaps from each design.

Coding intervals were derived from the GENCODE annotation (v49) [22], restricted to standard chromosomes and merged across transcripts to generate a non-redundant coding reference. Capture intervals were normalised, sorted and merged to produce non-overlapping genomic sets. Overlap between capture regions and the coding reference was computed using PyRanges. Coding coverage was defined as the proportion of coding bases overlapping capture regions, and capture specificity as the proportion of capture bases overlapping annotated exons. All base-pair calculations were performed on merged intervals to avoid double counting.

### Performance metrics

All metrics were computed within each workflow’s defined capture space to reduce bias introduced by differences in panel size and genomic breadth. However, this does not eliminate the influence of upstream differences in library preparation, input amount or deduplication strategy. Comparisons were performed using PyRanges for each pair and for all three platforms, identifying common and unique intervals, computing the total base-pair coverage. The resulting overlapping exome sizes are shown in Figure 1A.

**Figure 1.**
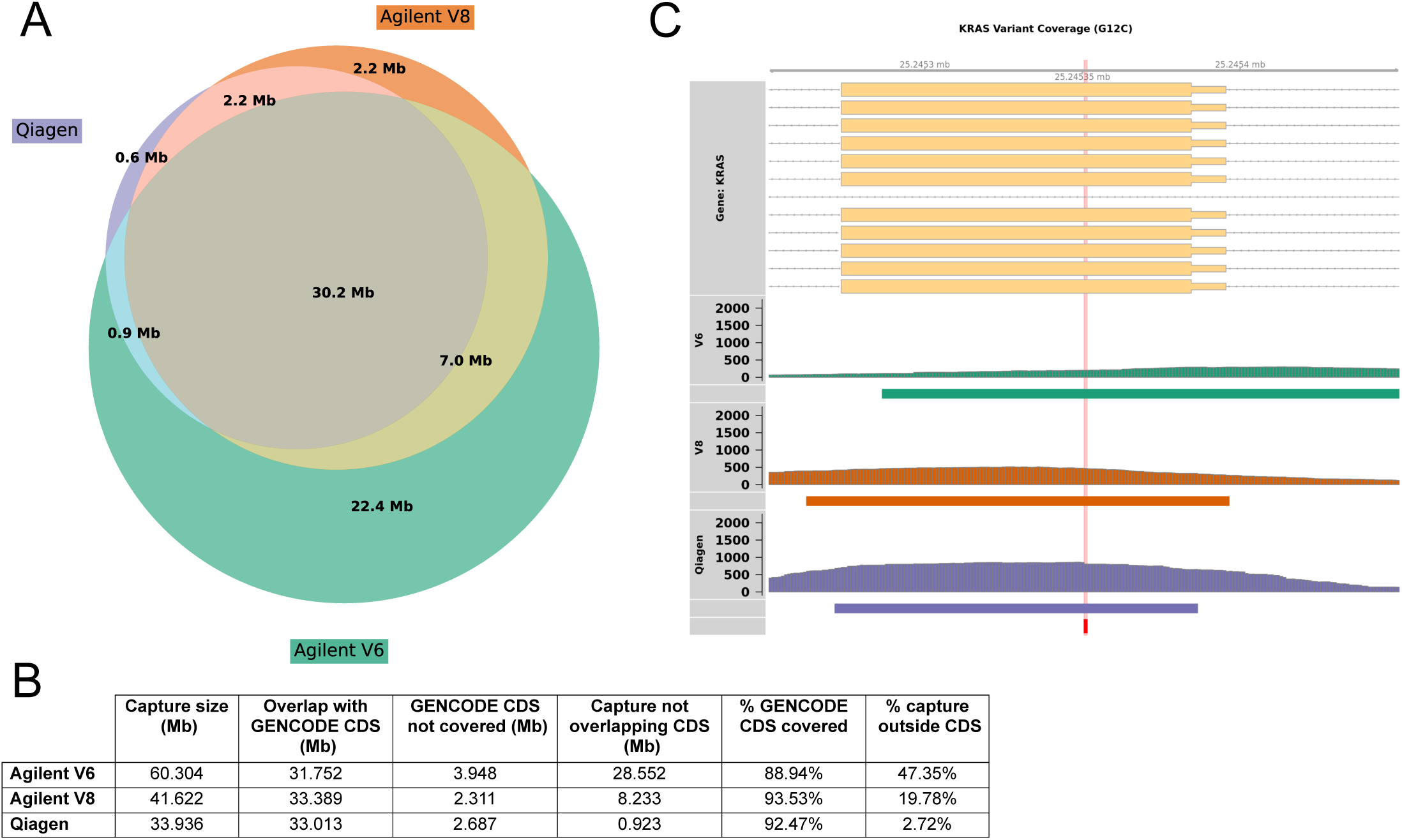
Comparison of exome capture regions of Agilent V6, Agilent V8 and Qiagen. **A**. Overlap of the reported capture regions for each whole exome capture chemistry tested, calculated from the available BED files. The consensus 30.2 Mb includes the majority of Qiagen and V8, which are tightly focused on CCDS. The largest panel Agilent V6 captures an additional 22.4 Mb unique bases, including probes for 5’- and 3’-UTRs. **B.** Exon coverage and capture specificity per chemistry, as compared to the GENCODE (v49) exome of 35.7 Mb. Capture sizes were derived from the *union* of all probes included the panel BED files **C.** IGV overview of probe coverage and typical sequencing depth at recurrent PDAC risk locus *KRAS* G12, for each chemistry. V8 and Qiagen are comparable with respect to coverage targeted on the exon, while the V6 design captures many more non-coding variants.

To compare the performance of the three WES panels, key quality control (QC) metrics were assessed. These metrics were derived from the raw, unmapped FASTQ file, the raw mapped BAM file and duplicate-marked BAM files (prior to UMI de-duplication) for each sample. For each kit, the analysis was restricted to the intervals specified from the manufacturer’s BED files.

For the specific short/long read average coverage calculations, the mapped pre-deduplicated BAM files were split on the following fragment sizes (0-150 bp, 150-300 bp, 300+ bp) using deeptools [23] alignmentSieve. BED files for each fragment size were generated using bedtools [24] intersect on the capture bed file and the sieved BAM file, before being sorted and merged using bedtools. Average coverage on the sieved BED files was computed using Genome Analysis Toolkit (GATK) [25] DepthOfCoverage.

### Mapping

For all three kits, raw sequencing reads in FASTQ format were trimmed using fastp [26] to remove adaptors. V6 and Qiagen data were mapped to the hg38 reference genome using BWA MEM (v0.7.17) [27]. The sorted BAM files were annotated with sample and platform identifiers and indexed using samtools (v1.19) [28]. Following alignment, PCR and optical duplicate artefacts were identified and marked using GATK (v4.6.2.0) [25] MarkDuplicates. Base Quality Score Recalibration (BQSR) was applied using GATK using a set of gold-standard SNPs and indels. This pipeline is detailed in Supplementary Figure S1A and was also used to obtain performance metrics for Agilent V8.

Since V8 data contains unique molecular indexes (UMIs), a separate pipeline was used for variant calling from this panel (Supplementary Figure S1B). Paired-end FASTQ files were aligned to the hg38 reference genome using BWA MEM with read group information specified. UMI sequences were extracted from read names and retained in the RX tag of the BAM file. Reads were query name-sorted, mate information corrected and grouped into molecular families using fgbio 1.0.0 (GroupReadsByUmi, paired strategy, zero mismatches) [29]. Single strand consensus and duplex consensus sequencing reads for V8 were generated, converted back to FASTQ using samtools, and realigned to hg38 to produce analysis-ready BAM files. Since UMI and non-UMI methods differ in error suppression, variant sensitivity in V8 reflects both exome capture design and downstream consensus processing strategy and should not be attributed to capture chemistry alone.

### Somatic variant calling and filtering

Variants were called on each sample for each capture method using samtools mpileup and VarScan (v2.3.9) [30] in consensus calling mode, and functionally annotated using RefGene [31], dbNSFP (v3.0a) [32], dbSNP (build 150) [33], COSMIC (v70) [34], gnomAD (v2.0.1) [35] and European 1000G (August 2015) [36] (Supplementary Figure S2A). The resulting multi-sample VCF files were filtered for variants present in dbSNP without COSMIC annotation, synonymous or unknown exonic variants, variants with population allele frequency >1% in gnomAD, recurrent artefacts, absence of alternate allele evidence in matched normal control samples, minimum sequencing depth ≥3 reads, alternate base quality ≥25, and strand support from both forward and reverse reads using a custom Python script and reference databases (Supplementary Figure S2B). To enhance recovery of clinically relevant PDAC variants, we incorporated a disease-focused gene whitelist based on the Agilent SureSelect targeted pancreas panel (Supplementary Table S3), permitting relaxation of the strand-support requirement for variants occurring in known risk genes. This rescue step was intended as a clinically motivated sensitivity analysis rather than a platform-neutral discovery metric.

Filtered variants per sample were lifted over to hg19/GRCh37 and analysed using the [Cancer Genome Interpreter](https://www.cancergenomeinterpreter.org/) (CGI) [37] to identify known (TIER 1) and predicted (TIER 2) driver mutations with potential therapeutic relevance.

### Copy number analysis

Copy-number analysis was performed on whole exome BAM files using samtools and analysed with ichorCNA (hg38) [38] using 1 Mb windows (minimum mapping quality 20), with GC/mappability correction, centromere masking, diploid ploidy assumption, and tumour purity estimation. Copy number profiles were examined at the *ERBB2* locus on chromosome 17 to assess evidence of amplification or gain, based on standard ichorCNA thresholds for log ratios, classifying outputs as either gain, amplification or high-level amplification. ERBB2 was selected because amplification had previously been identified in the matched V6 dataset [10], providing an opportunity to assess concordance across workflows.

## RESULTS & DISCUSSION

### Target Region Overlap

Capture footprint varied between workflows and was expected to influence both sequencing efficiency and downstream variant detection. Figure 1A illustrates the overlap of target regions across capture panels. Agilent V6 covers the largest genomic area, including 22.4 Mb of unique sequence and 30.2 Mb shared with Agilent V8 and Qiagen.

Each panel was evaluated against GENCODE (v49) [39] 35.7 Mb consensus coding sequence (CCDS) to quantify coding coverage and capture specificity (Figure 1B). Despite probing the largest capture area (60.3 Mb), Agilent V6 panel captured the lowest proportion of the CCDS (88.94%), compared with smaller, more targeted panels (from Agilent V8 (41.6Mb; 93.5%) and Qiagen (33.9 Mb; 92.5%)). Only 2.7% of the Qiagen panel targeted regions outside the CCDS, whereas fully 47.35% of V6 targeted non-exonic regions. While extensive non-coding coverage may be inefficient in detecting SNVs, it may provide additional opportunities for the identification of CNVs. These design differences are likely to affect sequencing efficiency, effective depth within clinically relevant regions and the trade-off between discovery breadth and actionable variant recovery. With this in mind, we investigated probe coverage and achieved sequence depth for each capture panel in mutational hotspots of key PDAC driver genes including *KRAS* (Figure 1C)*, TP53, SMAD4* and *CDKN2A* [40] (Supplementary Figure S3; Supplementary Table S4). V8 and Qiagen are comparable with respect to coverage targeted tightly to exons, while the V6 design captures many more non-coding variants. However, six of ten hotspot regions in *CDKN2A* revealed clear differences in capture potential between panels (Supplementary Figure S3D) not seen in other genes tested, with Qiagen better targeting exonic regions and canonical transcripts than V6 or V8. *CDKN2A* (encoding p16 tumour suppressor) has a somatic mutation rate of ∼18.5% (Pancreas Genome Phenome Atlas, GENIE dataset) [41, 42]. Its inactivation alongside common *KRAS* dysregulation promotes PDAC tumourigenesis [43]. Reduced probe coverage of *CDKN2A* has been shown to limit detection of key copy number changes by NGS, particularly in samples with low tumour content [44], which may influence panel selection and interpretation of results.

### QC Metrics

cfDNA is fundamentally different from intact genomic DNA extracted from cells or tissue, and these biological differences influence many of the QC metrics assessed, even before panel-specific effects are considered. Key factors include low input DNA amounts; short, unevenly fragmented DNAs; low tumour fraction; further signal dilution with non-tumour cfDNA, and the need for much deeper sequencing to compensate for these issues. These factors effectively limit the performance that could be expected from each kit in the context of low input cfDNA analysis.

### Depth and coverage

Across almost all sample/chemistry combinations, Agilent V8 generally showed higher coverage uniformity at sequencing depths >100X compared with V6 or Qiagen (Supplementary Table S5) within this dataset, despite Qiagen achieving higher raw sequencing depth (median range ∼650X-750X vs ∼350X-550X and ∼550X-700X, respectively), indicating that effective coverage in low-input ctDNA depends on capture efficiency and molecule recovery rather than sequencing depth alone. Indeed, Qiagen uniformity at >100X is consistently lower than either V6 or V8 in two of the three samples tested (∼83.3% vs 92.1%). This accords with a recent study showing V8 had excellent coverage statistics compared with three other exome enrichment kits from Roche, Vazyme and Nanodgmbio [45] and Agilent V7 [46]. However, cfDNA is naturally highly fragmented with end motifs and length distributions different from gDNAs, and distinct between tumour and normal plasma samples [47]. This can impact genomic representation from the start and contributes to uneven coverage (capture) [47] regardless of panel quality.

### On-target performance

Both Agilent platforms exhibited higher on-target performance across all samples, with on-target rates of >35% for V6 and V8, compared to a maximum of 33.9% for Qiagen (Supplementary Table S5). This suggests greater on-target efficiency for the V8 chemistry in capturing the intended genomic regions under target molecule scarcity, thereby maximising the proportion of useful sequencing reads. When normalised against requested sequencing depth, V8 appears to achieve more efficient capture and sequencing yield. These values are comparatively low for WES, where typical on-target rates are ∼50-70%, depending on capture chemistry, library insert size and DNA sample quality and quantity [48]. However, we should expect such reduced on-target rates for cfDNA, since shorter fragments hybridise less efficiently to the hybrid-capture probes used in WES, which require overlapping sequences to be most effective, i.e. these rates are likely exaggerated by cfDNA biology and low input cfDNA amounts in this experiment.

### Duplication rate

Duplication increases when capture efficiency is low and is almost unavoidable in low input cfDNA where library complexity is intrinsically reduced. Library complexity reflects the diversity of unique DNA fragments in a sequencing sample and determines the extent to which additional sequencing adds new (non-duplicate) reads. PCR duplication rates varied considerably between the three kits tested (Supplementary Table S5), with the lowest cfDNA input reactions (Qiagen; V8 < 5 ng; Supplementary Table S1) generally exhibiting much higher rates of duplication than for V6 reactions (>63.2% vs 39.9-46.4%) that included significantly more input (>25 ng). The data from V8 probe set neatly illustrate this, with P095_D (11.5 ng input) showing much lower duplication (34.6%) compared to either P045_A (63.2%) or P095_A (67.3%), where hybridisation conditions were similar. In the context of low input cfDNAs, high rates of duplication therefore reflect limitations in unique molecule availability and workflow-level efficiency of molecule recovery, rather than PCR artefacts alone. This can limit variant calling sensitivity, unless UMI-based consensus strategies are employed. This is a recognised challenge in cfDNA WES, particularly in cancer contexts where the tumour fraction is low [49]. These constraints highlight challenges for the application of cfDNA WES in clinical research settings, where detection of rare and/or therapeutically actionable variants is essential, and help explain why smaller targeted panels remain attractive for certain use cases.

### Adapter contamination

Qiagen showed markedly higher adapter contamination (17.58-25.04%) across all samples compared to either Agilent panel (9.95-15.01%) (Supplementary Table S5). Generally, adapter contamination may arise from inefficient ligation, sub-optimal library cleanup, or chemistry-specific characteristics [50]. However, it is of particular concern in cfDNA analysis, where a proportion of fragment sizes may be shorter than the read length (e.g. 2×150bp), although mean insert sizes across all samples and chemistries appear to be broadly consistent in our data: ∼171-189 bp. cfDNA proportions are also consistently high across all panels tested (74.78%-83.41%), suggesting that these results reflect genuine cfDNA signal, not fragmentation artefacts.

Limited input cfDNA (unique molecules) can also result in higher adapter:insert ratios, further exacerbating the problem. Here, starting amounts for Qiagen reactions were indeed lower than for V8 (Supplementary Table S1) in two of three samples, but these were not in proportion to the 1.7X-2.0X differences noted in adapter contamination.

Collectively, these results suggest that the Agilent V6 and V8 library preparation technologies outperform Qiagen across multiple quality control metrics in this cfDNA context. All three capture panels yield high quality data (Q30 89.3-93.1%) that maps extremely well (>99.9%) (Supplementary Table S5), so observed differences may be ascribed to sample features, library preparation and capture chemistry, not sequencing.

V6 provided the most consistent on-target rates, while V8 generally achieved the highest coverage and uniformity. By contrast, Qiagen demonstrated elevated duplication rates, increased adapter contamination, and lower uniformity, potentially compromising sequencing efficiency and variant calling accuracy. These findings highlight the reliability and efficiency of Agilent platforms for generating high quality WES data from challenging plasma cfDNA samples, even with lower than recommended input amounts.

### Somatic variant detection

In this context of low input (low mutational fraction), cfDNA variant calling poses a detection problem even with high sequencing depth, which can amplify strand-specific artefacts and generate false positives unless carefully filtered out [51, 52]. While strand bias filters were applied globally (Methods), given known locus-specific strand imbalance in cfDNA, particularly in established tumour drivers due to site-specific biological fragmentation patterns rather than technical artefacts [53, 54], strand balance requirements were relaxed for established PDAC risk genes (Supplementary Table S3; Agilent Targeted Pancreas Cancer assay) to enhance detection sensitivity for biologically relevant mutations, if present.

We compared the overlaps between the somatic variants called on samples analysed using the V6, V8 and Qiagen exome capture chemistries, after stringent filtering and whitelist rescue (Figure 2). Agilent V6 yielded the highest number of retained variants in each sample, largely consistent with its broader capture footprint: ∼3X-7X and ∼12X-18X as V8 or Qiagen respectively, disproportionate to the relative capture panel sizes of 60 Mb, 35.1Mb and 33.9 Mb (Supplementary Table S2), and not correlated with original starting amounts of cfDNA per reaction (Supplementary Table S1). Qiagen consistently captured the fewest variants in each sample. Both V8 and Qiagen panels also appear to capture many variants unique to each. Concordance between retained variant sets across all three workflows was low, with only 7-35 shared variants per sample, reflecting differences in bait footprint, effective depth, error suppression and filtering rather than capture design alone. Pairwise comparisons revealed that broadly, V6 and V8 shared the greatest number of variants across samples (67-212), while V8-Qiagen consistently showed the lowest overlap (79-131) (Figure 2A-C).

**Figure 2.**
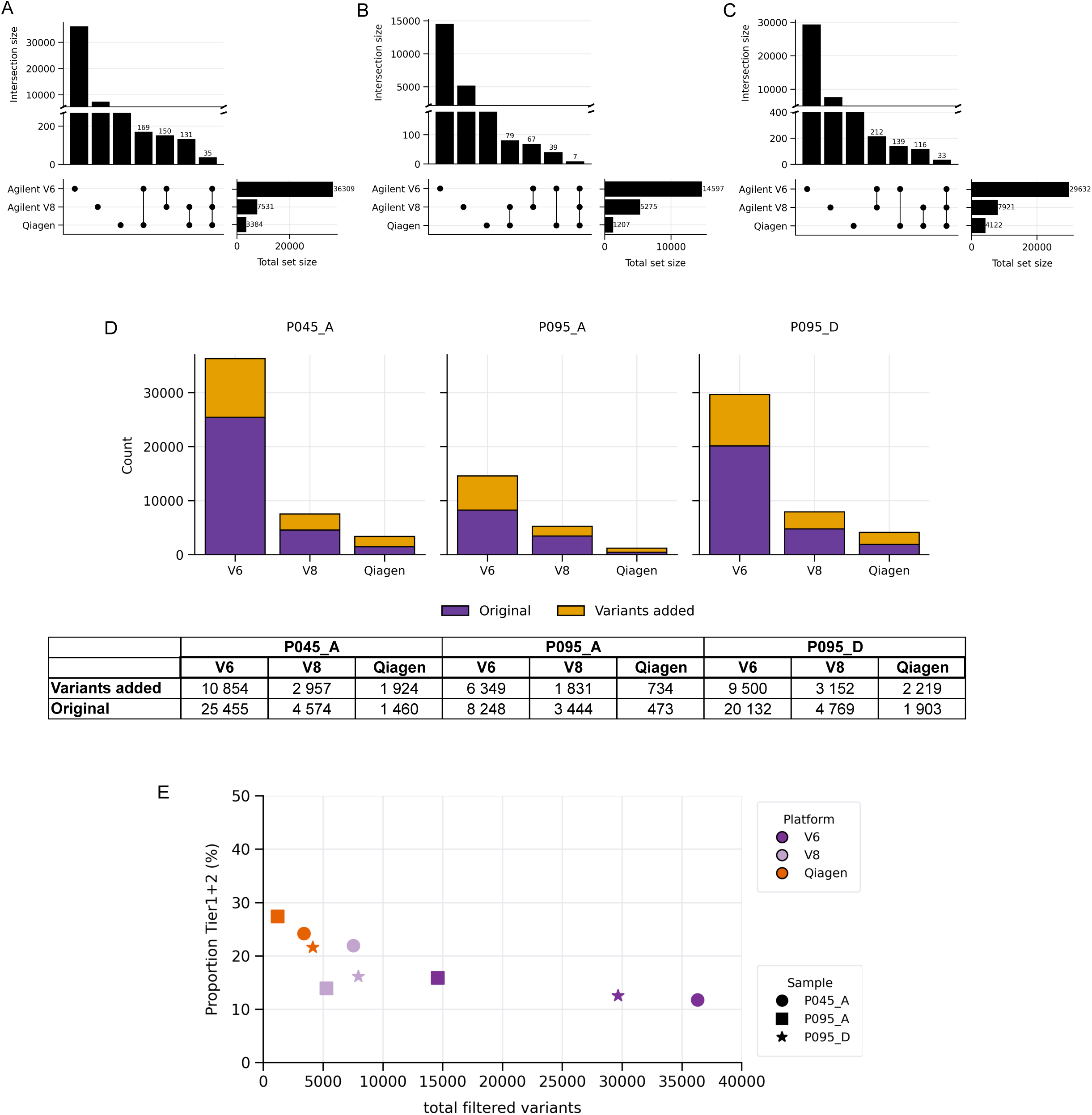
Unique and common variants identified in each sample by each capture chemistry. UpSet plots show the number of filtered somatic variants identified in samples **A.** P045_A, **B.** P095_A and **C.** P095_D by Agilent V6 (60 Mb panel), Agilent V8 (35.1 Mb panel) and Qiagen (33.9Mb) whole exome capture chemistries. Intersection sizes are shown on a linear scale with broken y-axis for clarity. Numeric labels show intersection sizes for shared probe sets, where intersections are ordered by size within each panel. **D.** The impact on variant call numbers per capture panel with whitelist filtering. **E**. Relative enrichment of actionable variants identified by CGI per capture chemistry.

Whitelist rescue contributed substantially to variant call numbers across all panels, particularly in V6 (6,349-10,854), with lower absolute recovery in V8 (1,831-3,152) and Qiagen (734-2,219) (Figure 2D). Overall, Qiagen identified the fewest SNVs, suggesting increased stringency but potentially reduced sensitivity. The significant impact of whitelist rescue rates >100% for all three samples support this: relaxing filtering stringencies for established risk genes for Qiagen’s highly targeted capture results in the retention of proportionally more disease-relevant variants. At the same time, the much higher variant counts in V6 reflect its substantial genomic breadth; with relaxed fragment constraints this likely includes a higher proportion of lower confidence calls in these low tumour fraction samples. Functional annotation of the final filtered variants per patient in CGI supports this, with ∼22-28% Qiagen variants identified as either TIER 1 (known) or TIER 2 (predicted) oncogenic drivers, compared to ∼12-16% of Agilent V6 variants (Figure 2E; Supplementary Table S6). Variant yield should therefore be interpreted as a conditional output of capture breadth, molecule recovery, consensus strategy and filtering behaviour, rather than as a direct estimate of true biological sensitivity. Indeed, as others have shown [55–57], pipeline design and choice of filtering strategy can substantially affect the number of variants retained (Supplementary Figure S4).

### Tumour-Plasma Concordance

Supplementary Figure S5 shows the overlap of filtered variants between tumour and matched patient plasma samples. Overall concordance was low (0.12-0.47%) across all samples and platform as expected in low tumour fraction PDAC cfDNA, although this was reported to increase to >55% when multiple serial plasma samples were combined in an earlier study using only V6 data [10] and a tumour-informed variant filtering approach (Supplementary Figure S4). Except for Qiagen P45_A and P95_A, plasma samples consistently yielded more filtered variants than tumour tissue, reflecting both the greater ability of plasma samples to capture tumour heterogeneity beyond tumour biopsies alone, and the increased background signal associated with low input, low tumour fraction cfDNA [10, 49]. V6 detected the fewest tumour variants, consistent with its less targeted capture region, and, likely as a result, achieved the lowest mean tumour-plasma concordance (0.17% compared to 0.35% for V8 and 0.44% for Qiagen). The consistently larger tumour-plasma overlap across Qiagen samples is likely due to its smaller capture region tightly targeting exons and canonical transcripts (Supplementary Figure S3), increasing the likelihood of shared variants between tumour and plasma. Higher concordance may therefore reflect both improved specificity and reduced variant yield.

### Copy Number analysis

Analysis of both Qiagen and V8 data using ichorCNA [38] reproduced the *ERBB2* copy number amplification signal previously identified in longitudinal V6 data [10], demonstrating moderate gain at the locus (log2 ratio 0.206) (Figure 3) and supporting the feasibility of recovering concordant focal CNV information across workflows in at least some cases. *ERBB2* amplifications are present in ∼2% of PDACs [58]; targeting this subset of patients with anti-HER2 therapies has been reported, with mixed results [58, 59]. Resistance to HER2-targeted therapies has been observed in PDAC patients with activating *KRAS* mutations [58] as well as other solid tumours [60].

**Figure 3.**
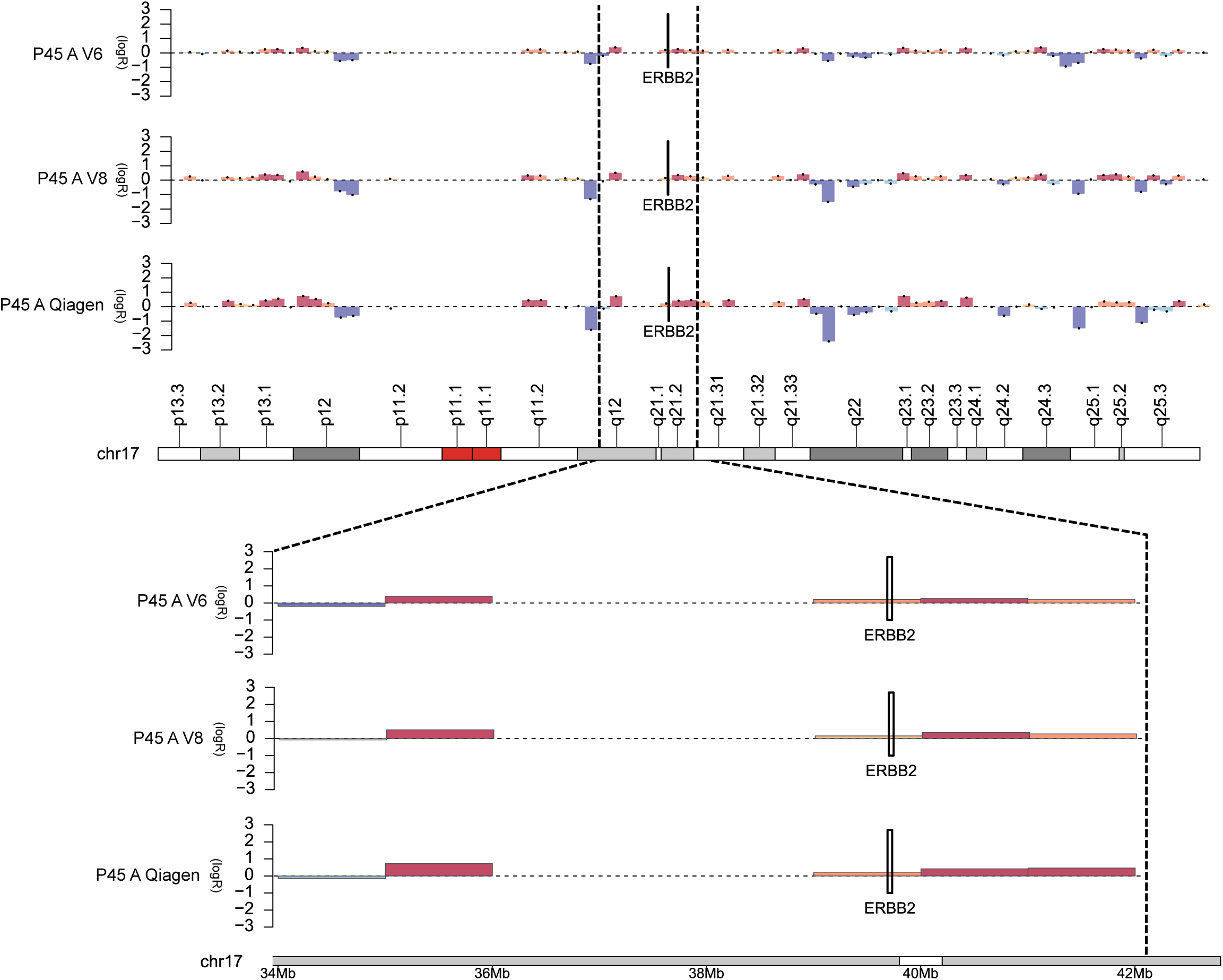
CNV detection by exome capture chemistry. Copy number profiles for chromosome 17 in sample P045_A generated using three capture chemistries (V6, V8, Qiagen) are shown. Log ₂ copy number ratios are plotted along the y-axis, with genomic position (Mb) along the x -axis. Regions of copy number gain and loss are indicated in red and blue, respectively, with segmented copy number estimates overlaid. The *ERBB2* locus is marked by vertical dashed lines in the genome-wide view (top). The lower tracks show a magnified view of the *ERBB2* region (∼34 - 42 Mb), demonstrating consistent focal amplification across all platforms. Cytoband structure is indicated along the chromosome ideogram.

More recent tools such as SavyCNV [61] and OFF-PEAK [62] utilise off-target reads in exome sequencing by treating them as low coverage, genome wide read depth signals, and use these to identify large deletions or duplications that otherwise would not be detectable in exome data. These approaches depend on capture size and off-target coverage [61], suggesting that streamlined panels with limited non-coding representation like Qiagen (2.72%; Figure 2B) may be less informative for off-target-based CNV inference, whereas almost half of the V6 panel targets regions *outside* the CCDS (47.35%; Figure 2B).

### Unique Molecular Indexes

UMIs (molecular barcodes) are short, random nucleotide sequences in the NGS reaction adapters, which are ligated to cfDNA fragments before PCR amplification, giving each starting molecule a unique tag. These permit PCR duplicates to be identified and support error correction and accurate quantification, which is particularly relevant to low input cfDNAs with high sequencing depth.

After strand bias filtering, single strand consensus sequencing (SSCS) reads from Agilent V8 retained 3,444-4,769 variants across samples, compared to 119-203 for duplex consensus sequencing (DCS) (Supplementary Table S7-UMIs). Following whitelist rescue, final variant counts increased by ∼20% for DCS to 143-239 and ∼60% for SSCS to 5,275-7,921. DCS consistently yielded substantially fewer final variant calls than SSCS, reflecting its increased stringency and suppression of sequencing artefacts through the requirement for complementary strand support. However, this stringency has important implications for ctDNA analysis in low abundance states (e.g. early-stage disease) or tumour types (e.g. PDAC).

PDAC releases limited tumour-derived DNA into circulation, reducing the likelihood of recovering both strands of an original fragment. Consequently, true variants may only be represented in single-strand consensus families and fail to achieve duplex support. The markedly reduced variant counts observed with DCS suggest that such variants are likely excluded. In contrast, SSCS retained substantially more variants after filtering, indicating improved sensitivity for low-abundance variants. Although this may include more false positives, it better reflects the biological constraints of PDAC cfDNA, where detection is inhibited by low tumour fraction.

Overall, these findings highlight a trade-off between sensitivity and specificity, suggesting that despite its increased false-positive risk, SSCS analysis involving UMIs (Agilent V8) may be more appropriate for variant detection in low-shedding tumours such as PDAC, or in detection of early stage or minimal residual disease.

### WES vs targeted gene panels

Since gene panels offer more sensitive (higher depth) sequencing of curated risk loci, often with lower sample requirements [17] and perform well for hotspot SNVs and known oncogenic substitutions, we compared our WES-derived findings with what may have been identified by screening the samples using off-the-shelf gene panels for somatic variant detection. Several assays are available from each provider; we focused on the most comprehensive and/or those designed specifically for cfDNA analysis, and for which the BED files were freely available. We included Agilent (SureSelect Cancer CGP panel; exons of 679 genes; min. input 50 ng), Qiagen (QIASeq targeted DNA human comprehensive cancer panel; exons of 275 genes; min. input 10 ng) and Agilent SureSelect Targeted Cancer (Pancreas) assay (exons of 58 genes, a subset of their CGP assay; min. input 10 ng) for comparison. To our knowledge, the latter is currently the only commercially available pancreas-specific gene panel (Supplementary Table S3).

Comparison of the WES-derived variant sets against the panel capture BED files showed strong chemistry-dependent differences in WES variant yield and theoretical panel coverage (Supplementary Table S8). The QIAseq panel consistently overlapped with ∼1% of WES-identified variants across samples and chemistries, reflecting its highly targeted design for capturing a refined subset of tumour-associated coding variants, distinct from the broader set of somatic variants detectable by WES: the low percentages are consistent even as WES variant yield reduces with newer chemistries from ∼36K to ∼1,200 variants.

By contrast, the Agilent Pancreas Tumour panel consistently captures ∼85%-95% of the CGP-overlapping WES variants across all three samples and chemistries, suggesting that for pancreas-specific driver discovery alone, the added breadth the CGP offers is modest. Considering particularly the Agilent Pancreas panel overlap with WES captures, this increases markedly as newer capture chemistries and more stringent filtering reduces total WES variant burden (V6: 29.14%-42.29%; V8: 33.06%-55.0%; Qiagen 50.95%-57.17%), their effective variant outputs converge on the higher confidence, disease-associated variants that targeted panels are optimised to detect where they are ideal for screening in high risk populations. WES remains essential for unbiased discovery and exploratory analyses of early genomic changes, and identification of composite, complex biomarkers like TMB, MSI, or pathway-level changes like HRD or mismatch repair deficiency and which are associated with response to immune checkpoint inhibitors in various solid tumours [63] including pancreatic cancer [18, 19].

## Discussion & Conclusion

We evaluated the sensitivity, accuracy and potential clinical relevance of WES of plasma ctDNAs in a realistic context of limited sample availability, by systematically and quantitatively comparing the performance of competing exome capture chemistries designed to accommodate low amounts of input cfDNAs. We benchmarked these existing data from a well-characterised cohort of PDAC patients [10].

This analysis demonstrates clear performance differences between the three WES library preparation technologies when applied to cfDNA samples. The Agilent V6 and V8 kits generally outperformed the Qiagen kit in the critical areas of coverage uniformity, depth, and on-target specificity. This suggests that the Agilent capture technologies are more efficient at enriching for the targeted exomic regions in these challenging sample types (low cfDNA input amounts), where baseline capture efficiency is input-limited, not kit limited.

A key differentiator was the PCR duplication rate, a crucial indicator of library complexity. The substantially higher duplication rates seen with the Qiagen kit indicate a lower initial library complexity, meaning more sequencing reads were required to achieve the same effective coverage. The V8 kit showed the lowest duplication rates, suggesting it is the most efficient of the three at capturing and converting unique DNA fragments into sequenceable library molecules.

We acknowledge that our findings are based on a small sample set, limited to patients for which we had data for all three chemistries tested. This limits the generalisability of our results to other tumour types or disease stage with higher cfDNA availability or higher tumour-derived fractions, including advanced diseases and molecular subtypes associated with increased ctDNA shedding. Also, although DNA-level batch effects are minimal compared to RNA-based analyses, library preparation, sequencing platform and operator differences may all impact data generation. However, these data are still informative in this context of longitudinal patient sampling of low tumour fraction plasma with realistically low sample availability. Importantly, this study is intended as a technical and descriptive benchmarking exercise. It does not establish absolute limits of detection, sensitivity or specificity, which would require independently validated reference materials sequenced across platforms.

While all three kits produce usable sequencing data, the Agilent V6 and V8 technologies demonstrate superior performance for low input cfDNA WES applications. They provide deeper, more uniform coverage with a higher on-target rate and a more efficient use of sequencing resources, making them a more suitable choice for studies where maximising the recovery of genome-wide information from limited-input samples is critical.

**Supplementary Table S1.** cfDNA input amounts per sample, per chemistry tested.

|  | **Agilent v6** | **Agilent v8** | **Qiagen** |
| --- | --- | --- | --- |
| **Input plasma volume/sample** | ~ 3ml | ~500 ul | ~500 ul |
| **Patient ID** | **Input/library reaction (ng)** |  |  |
| P045_A | 25 | 2.74 | 3.3 |
| P045_C | 50 | - | 3.5 |
| P045_D | 90 | - | - |
| P045_E | 15.69 | 19.1 | - |
| P095_A | 7.7 | 3.2 | 2.02 |
| P095_B | 13.5 | 1.03 | - |
| P095_C | 14 | - | - |
| P095_D | 33.8 | 11.52 | 7.02 |

Only samples with data available for all three chemistries were included in this report: P045\_A, P095\_A, P095\_D.

**Supplementary Table S2.** Experimental details for each of the chemistries tested.

|  | **Agilent v6** | **Agilent v8** | **Qiagen** |
| --- | --- | --- | --- |
| **Library preparation** | ThruPLEX<br>PlasmaSeq | SureSelect XT<br>HS2 DNA | QIAseq Ultralow<br>Input |
| **Listed minimum input required** | 1 ng | 10 ng | 10 pg |
| **Total capture size** | 60 Mb | 35.1 Mb | 33.9 Mb |
| **Sequences targeted** | RefSeq<br>CCDS<br>GENCODE<br>HGMD<br>5'- and 3'-UTRs | RefSeq<br>CCDS<br>GENCODE<br>TERT promoter | CCDS |
| **Blocking oligonucleotides** | i5 + i7 xGen<br>universal blocking<br>oligos (IDT) | Yes; included | One-4-All; included |
| **Probe type** | 120 nt biotinylated<br>cRNA | 120 nt biotinylated<br>cRNA | 120 nt biotinylated<br>dsDNA |
| **Probe hybridisation** | 65°C; 25h<br>(min. 24h) | 67.5°C; 21h<br>(min. 16h) | 60 °C; 18h<br>(min. 16h) |
| **UMIs** | no | Yes, included | No |
| **Post-capture amplification cycles (dependent on available library amount; ng)** | 10 | 10 | 8 |
| **Approx. target read depth** | ~500X | ~1300X | ~1100X |

Agilent v6: SureSelect XT2 human all exon v6, Cat. 5190-8872
Agilent v8: SureSelect XT human all exon v8, Cat. 5191-6879
Qiagen: Qiagen Human Exome, Cat. 333937
CCDS: consensus coding sequence
UMI: unique molecular index
All technical details are extracted from the respective User Manuals.

**Supplementary Table S3.** PDAC genes whitelist.xls.

**Supplementary Table S4.** PDAC hotspot seq depths.xls.

**Supplementary Table S5.** QC metrics.xls.

**Supplementary Table S6.** CGI variants.xls.

**Supplementary Table S7.** UMIs.xls.

**Supplementary Table S8.** Detection of WES variants by targeted sequencing panels across patients, timepoints, and platforms. Values represent the number of variants detected and percentage of total WES variants.

| Sample | Capture | WES<br>(n) | Agilent Pancreas<br>(58 genes)<br>n (%) | Agilent CGP<br>(679 genes)<br>n (%) | QIAseq<br>(275 genes)<br>n (%) |
| --- | --- | --- | --- | --- | --- |
| **P045_A** | V6 | 36,309 | 11,697 (32.22) | 10,579 (29.14) | 139 (0.38) |
|  | V8 | 5,093 | 2,871 (56.37) | 2,801 (55.00) | 26 (0.51) |
|  | Qiagen | 3,384 | 1,832 (54.14) | 1,777 (52.51) | 30 (0.89) |
| **P095_A** | V6 | 14,597 | 6,531 (44.74) | 6,173 (42.29) | 80 (0.55) |
|  | V8 | 5,275 | 1,894 (35.91) | 1,744 (33.06) | 32 (0.61) |
|  | Qiagen | 1,207 | 700 (58.00) | 690 (57.17) | 14 (1.16) |
| **P095_D** | V6 | 29,632 | 10,151 (34.26) | 9,263 (31.26) | 134 (0.45) |
|  | V8 | 7,921 | 3,170 (40.02) | 2,992 (37.77) | 36 (0.45) |
|  | Qiagen | 4,122 | 2,177 (52.81) | 2,100 (50.95) | 32 (0.78) |

\* Variant recovery was calculated by intersecting WES variant positions with panel BED regions. Variants falling within panel intervals were counted as captured, and proportions were computed relative to total WES variants per sample.

## Ethics approvals

Whole blood, plasma and tumour biopsy tissues from patients with histologically confirmed pancreatic ductal adenocarcinoma were obtained from the Barts Pancreatic Tissue Bank (www.bartspancreastissuebank.org.uk, Research Ethics Committee reference 13/SC/0592, renewed 18/SC/0629, renewed 23/SC/0282). All patients provided written, informed consent for the processing and use of their samples and publication of study results. All patient data included in this manuscript is non-identifiable.

## Availability of data

Agilent v6 sequence data has been deposited at the European Genome-phenome Archive (EGA) (https://ega-archive.org/datasets/) under accession number EGAD00001008593 (Study ID: EGAS00001005981; Data Access Committee: EGAC00001002556). Additional data will be submitted and available from the EGA upon acceptance.

## Author contributions

Conception and design: CC, HRA; patient sample and clinical data acquisition and curation; PCRFTB, HMK, CC: sample preparation & data acquisition: HRA, bioinformatic analysis and/or interpretation: LGEJ, GJT, CM, HRA; Writing – original draft and revision: HRA, LGEJ, CM, GJT; Writing – review and editing of final draft: All authors; funding acquisition: HMK, CC; Supervision: CC, HRA.

## Acknowledgements

We are grateful to all the patients and their families who have donated blood and tissues to the Barts Pancreas Tissue Bank (https://www.thepancreastissuebank.org/, and to the BPTB staff for establishing the framework for collection and distribution of samples and clinical data.

## List of abbreviations

CCDS: consensus coding sequence
cfDNA: cell-free DNA
CGP: comprehensive gene panel
CN: copy number
CNA: copy number alteration
CTC: circulating tumour cell
ctDNA: circulating tumour DNA
HRD: homologous recombination deficiency
MBC: molecular barcode
MSI: microsatellite instability
NGS: next generation sequencing
PDAC: pancreatic ductal adenocarcinoma
QC: Quality Control
SNV: single nucleotide variant
TEP: tumour-educated platelets
TMB: tumour mutational burden
UMI: unique molecular index
VAF: variant allele frequency
WES: whole exome sequencing

