## Supplementary material for "Systematic benchmarking of low-input whole exome sequencing workflows for longitudinal ctDNA profiling in pancreatic ductal adenocarcinoma": Suppl. Fig S2

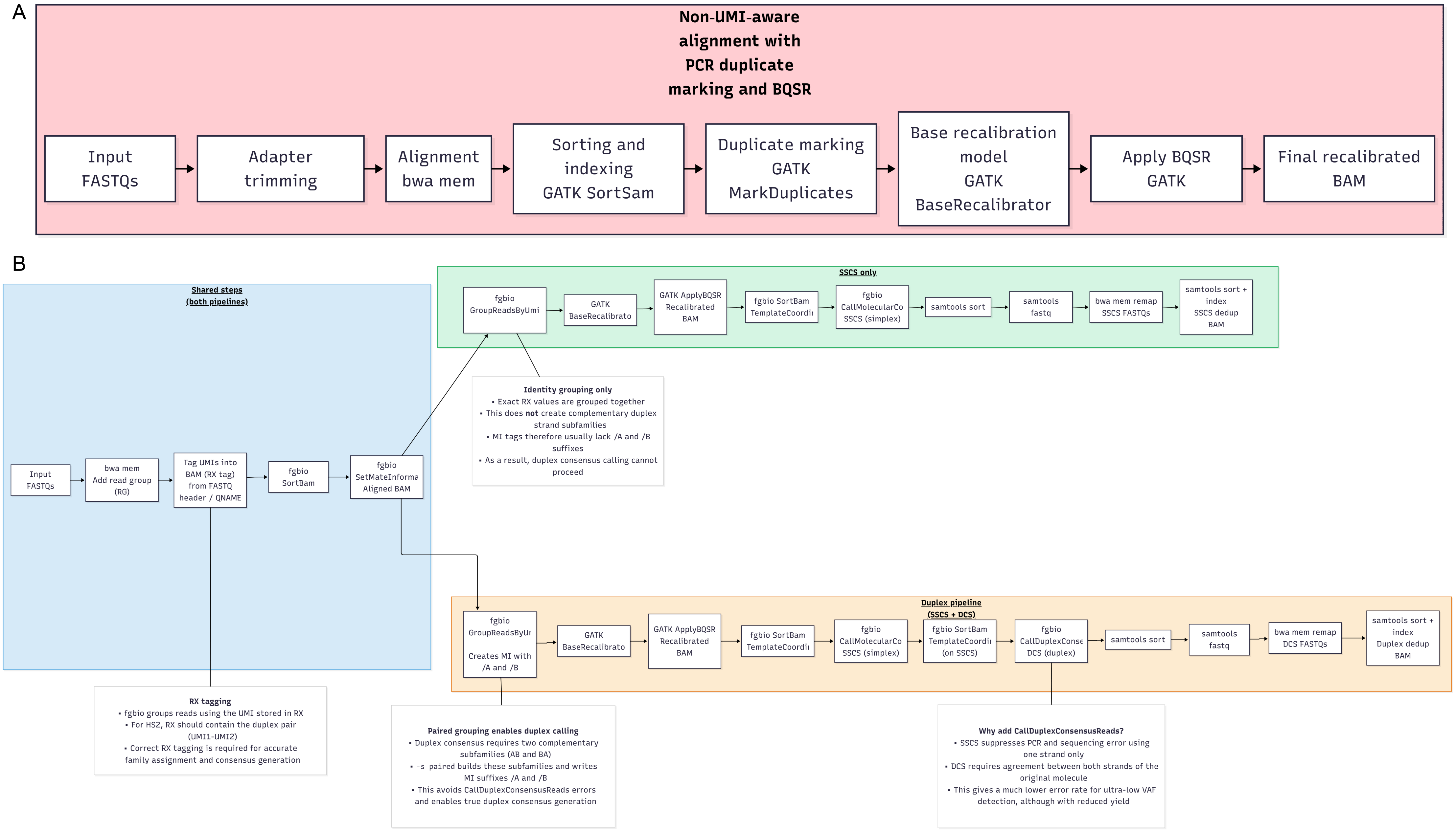


**Supplementary Figure S1.** Overview of mapping pipelines for data deduplication

**A**. Analytic pipeline for Agilent V6 and Qiagen data: non-UMI aware alignment with PCR duplicate marking and base quality score recalibration (BQSR).

**B**. Mapping pipeline for data generated from Agilent V8 data containing UMIs (molecular barcodes).
