## Supplementary material for "Systematic benchmarking of low-input whole exome sequencing workflows for longitudinal ctDNA profiling in pancreatic ductal adenocarcinoma": Suppl. Fig S2

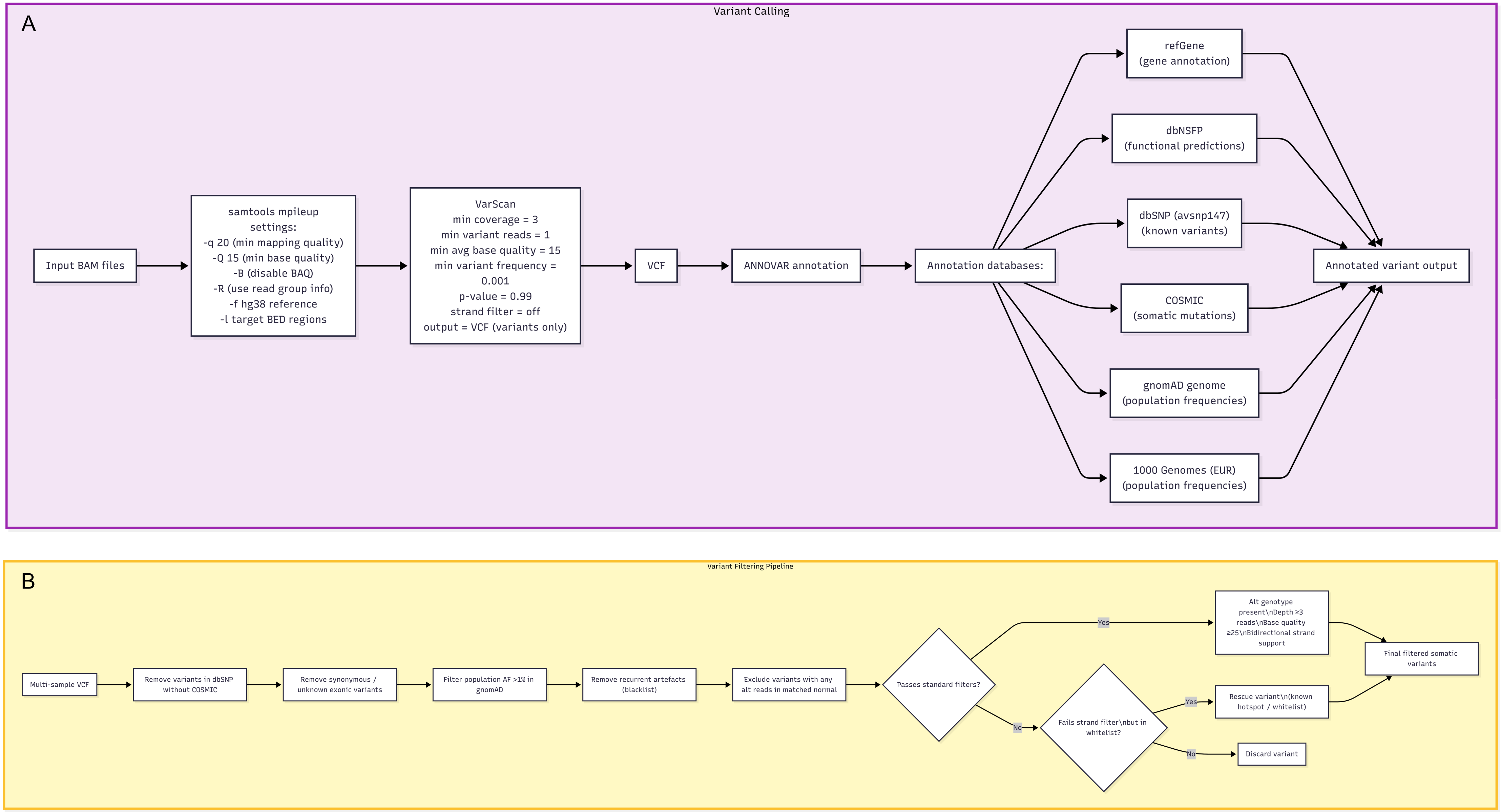


### Supplementary Figure S2

**A**. Variants were called on each sample for each capture method and annotated using various databases. **B**. Somatic variants were filtered to identify likely ctDNA mutations. 58 genes included in the Agilent SureSelect pancreas-specific cancer gene panel were used as a whitelist and incorporated with more permissive strand support for variants occurring in established pancreatic cancer risk genes.
