## Supplementary material for "Systematic benchmarking of low-input whole exome sequencing workflows for longitudinal ctDNA profiling in pancreatic ductal adenocarcinoma": Suppl. Fig S3

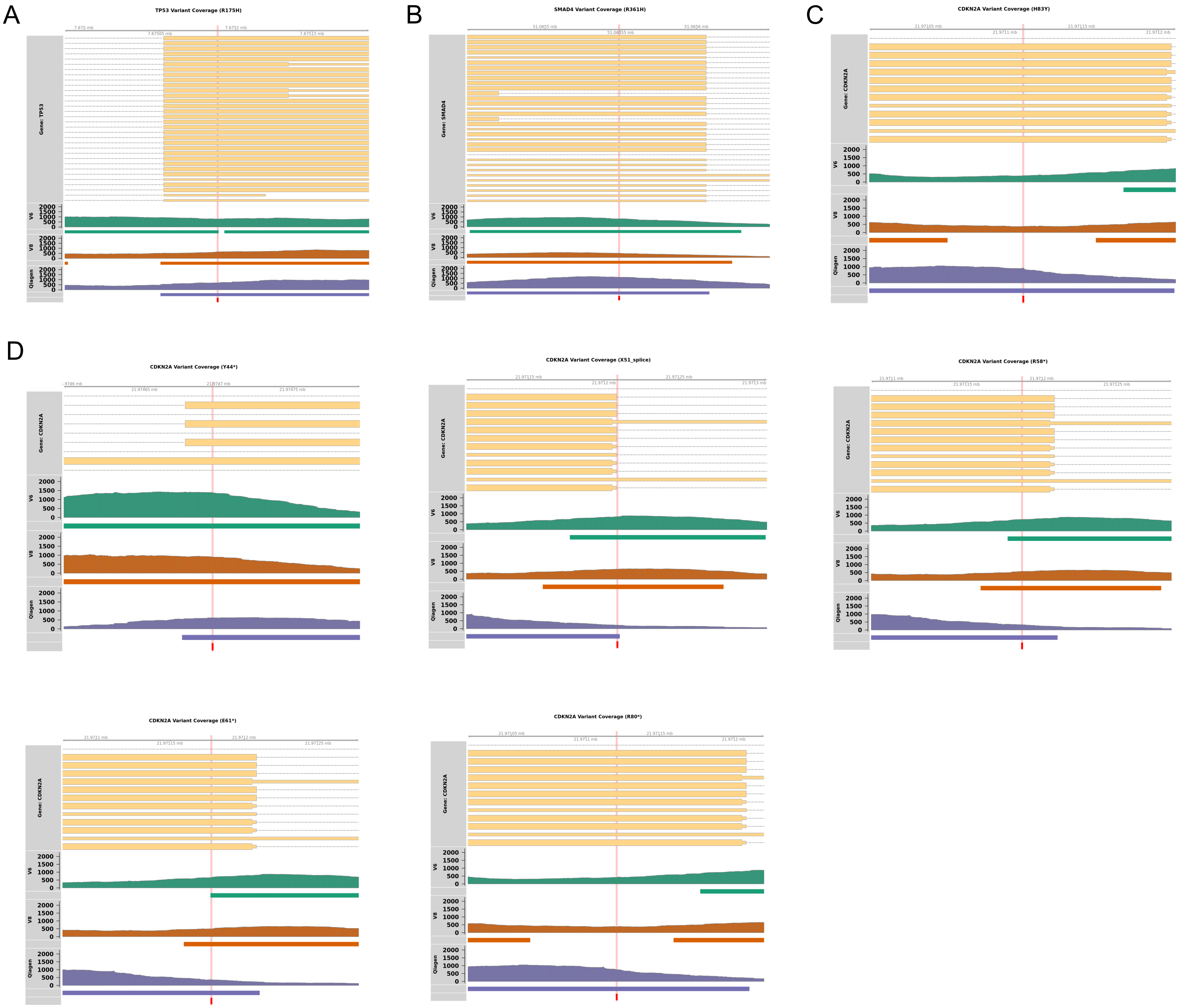


**Supplementary Figure S3**. Probe coverage and sequencing depth

Probe coverage and typical sequencing depth around recurrent risk loci regions in known PDAC driver genes, using sample P045_A as exemplar. **A.** *TP53* R175, **B**. *SMAD4* R361, **C**. *CDKN2A* H83. PDAC-specific representative amino acids were identified with Pancreas Genome Phenome Atlas (Oscanoa et al, 2025) in hotspot regions identified by Varghese et al (2025). **D**. Six of ten mutation hotspots in *CDKN2A* (Y44, X51, R58, E61, R80) revealed differences in exonic capture between panels.
