## Supplementary material for "Systematic benchmarking of low-input whole exome sequencing workflows for longitudinal ctDNA profiling in pancreatic ductal adenocarcinoma": Suppl. Fig S4

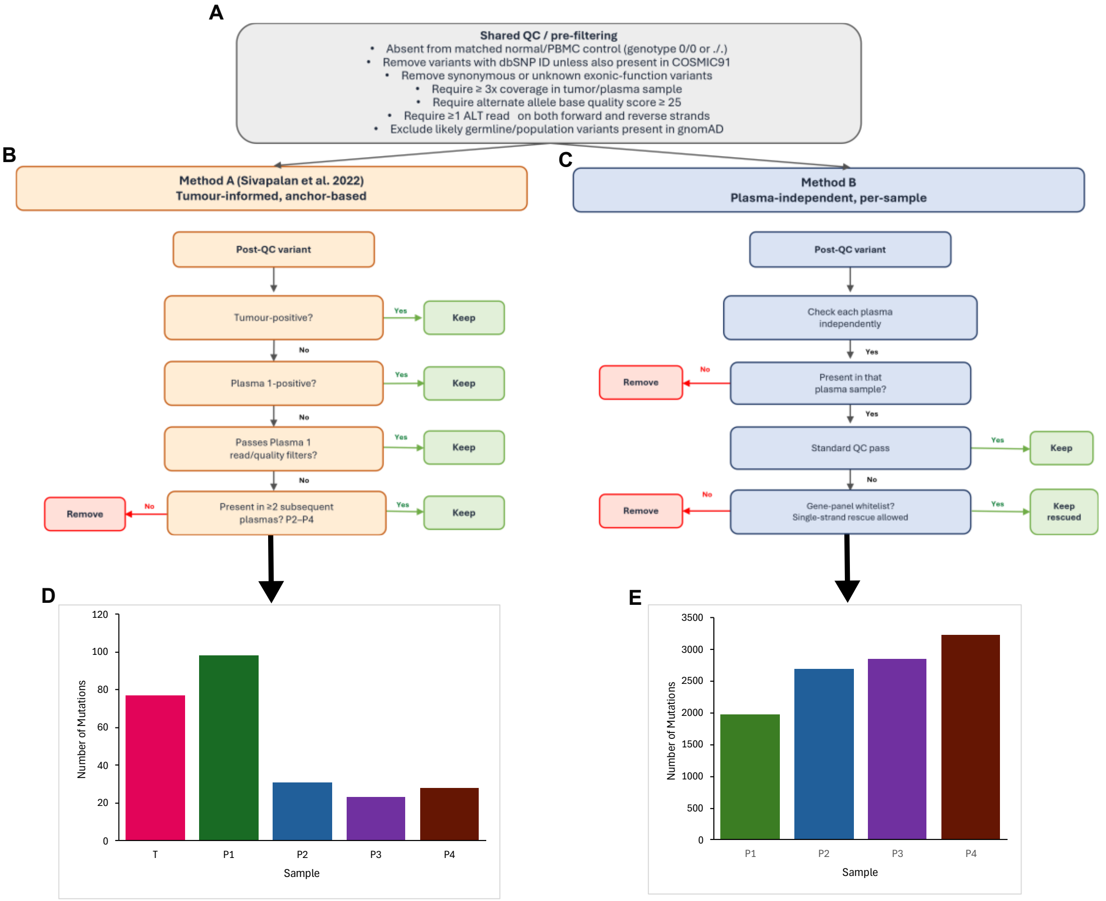


**Supplementary Figure S4 -** Comparison of tumour-informed and plasma-independent filtering strategies. The comparison was performed using the multi-sample VCF for patient P95 from *Sivapalan et al. 2022*. For the plasma-independent analysis, tumour and normal samples were removed from the input VCF so that filtering was applied only across the serial plasma samples. **A.** Shared QC and pre-filtering steps applied before method-specific filtering. **B.** Method A, based on *Sivapalan et al. 2022*, is tumour-informed and anchor-based. Variants are retained if detected in the tumour, or, if tumour-negative, if detected in Plasma 1 and passing Plasma 1 read-depth and quality filters. Variants absent from both tumour and Plasma 1 are retained only if present in at least two subsequent plasma samples. Variants detected in either the PBMC/normal control or the additional tissue-normal control are excluded before tumour/plasma prioritisation. **C.** Method B is plasma-independent and per-sample. Each plasma sample is filtered independently after QC, and variants are retained for any plasma timepoint in which they are detected. Variants failing standard QC are removed, except for gene-panel whitelist variants that may be rescued with single-strand support. **D-E.** Retained variant counts for Method A and Method B, respectively. These results show that pipeline design and variant-filtering strategy substantially affect retained variant counts, even when applied to the same input data.
