## Supplementary material for "Systematic benchmarking of low-input whole exome sequencing workflows for longitudinal ctDNA profiling in pancreatic ductal adenocarcinoma": Suppl. Fig S5

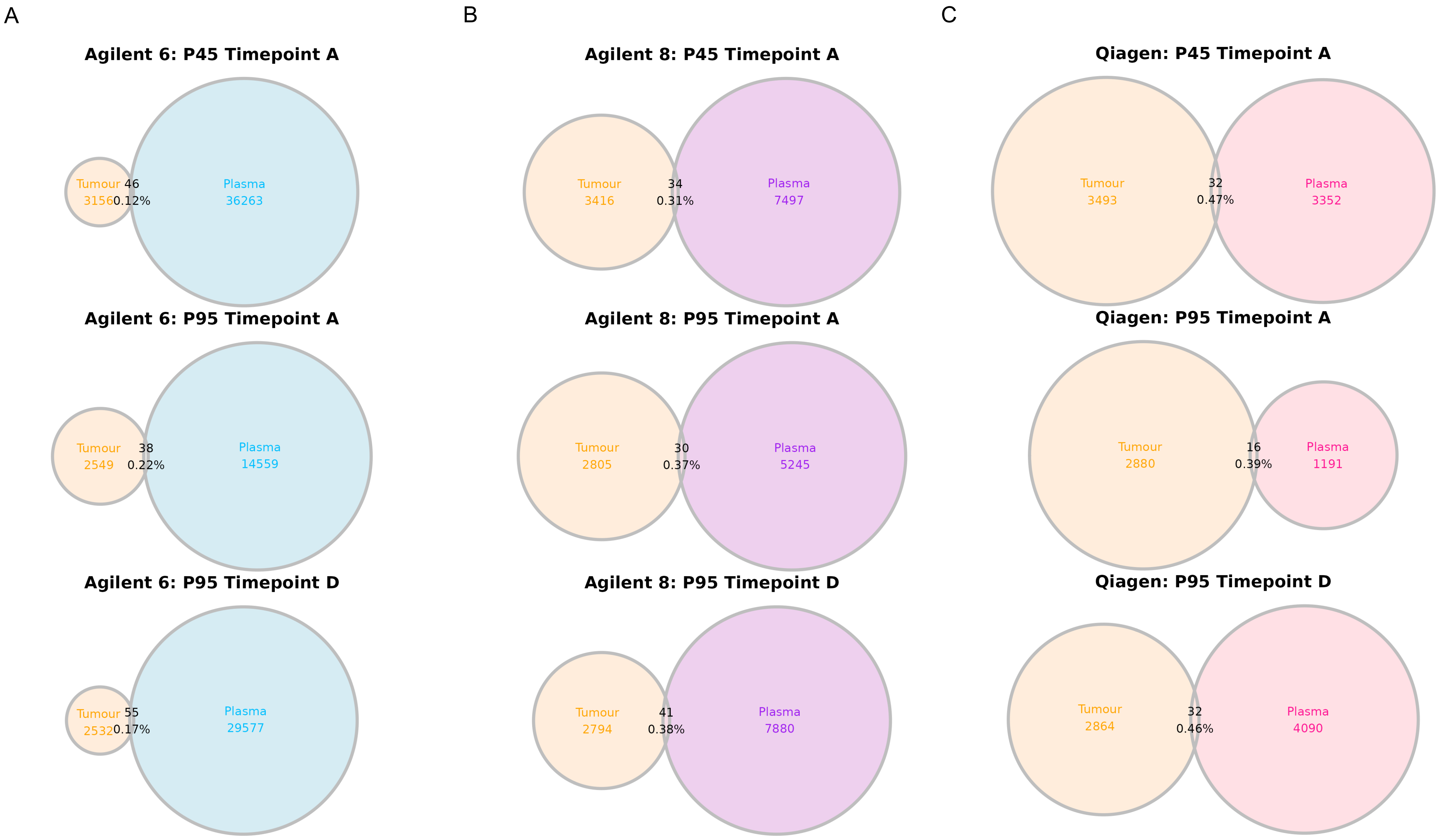


**Supplementary Figure S5.** Tumour vs plasma concordance

Overlap of filtered somatic variants between tumour tissue and matched patient plasma samples. Variants were matched by chromosome, start and end position, with percentages indicating proportion of total variants shared between tumour and each individual plasma sample, . captured by **A.** Agilent V6, **B.** Agilent V8 and **C**. Qiagen whole exome panels.
